# Ultra-deep duplex sequencing reveals unique features of somatic evolution in the normal tissues of a family with Li-Fraumeni syndrome

**DOI:** 10.64898/2026.01.12.699071

**Authors:** Hunter L. Colegrove, Marianne E Dubard-Gault, Henry Marshall, Brendan F. Kohrn, Thomas H. Smith, Zachary K. Norgaard, Fang Yin Lo, Elizabeth K. Schmidt, Jacob E. Higgins, Charles C. Valentine, Desiree A. Marshall, John I. Clark, Eric Q. Konnick, Jesse J. Salk, Marshall S. Horwitz, Raheleh Rahbari, Alison F. Feder, Rosa Ana Risques

## Abstract

Li-Fraumeni Syndrome (LFS) is caused by germline pathogenic variants in *TP53* which predispose carriers to early onset cancer across multiple tissues. While genomically profiling those cancers has revealed factors contributing to their formation, little is understood about how LFS impacts clonal evolution in healthy tissues preceding cancer. Here, we use ultra-deep duplex sequencing (mean ∼15,000× depth) to investigate somatic mutation and selection in a family carrying the germline *TP53* p.R181H pathogenic variant and a cohort of non-carrier controls. In blood samples, the germline variant is associated with more mutations in a panel designed to capture genomewide mutagenesis, and with reduced positive selection on somatic *TP53* mutations, despite confounding by chemotherapy treatment in one individual. *DNMT3A* and *TET2* mutations appear positively selected and *GATA2* mutations negatively selected across the cohort, independent of the p.R181H status. Extensive multi-tissue sampling of 22 non-cancerous and 6 cancerous samples was also performed at autopsy in one individual with LFS who succumbed to esophageal cancer. Cross-tissue analysis reveals excess mutations in sun-exposed skin, esophagus and chronically-inflamed stomach tissue, and concordant mutations in the p.R248 hotspot of *TP53* across most (18/28) tissue samples. Most somatic *TP53* mutations in LFS that can be assessed for phase arose on the chromosomal copy lacking the p.R181H variant. Our study reveals how the germline p.R181H variant reshapes baseline somatic mutation and selection in normal tissues and highlights the importance of understanding early somatic evolution in LFS prior to cancer development and treatment.

## Introduction

Li-Fraumeni syndrome (LFS) is a cancer predisposition syndrome caused by an inherited or de novo germline pathogenic variant in *TP53*. Individuals with LFS are at risk of developing multiple cancer types starting early in life, with an up to 24 times higher incidence of cancer than the general population (Mai et al. 2016; de Andrade et al. 2021; McBride et al. 2014). The increased cancer risk is due to the higher likelihood of completely inactivating *TP53* via a second-hit mutation or loss of heterozygosity, given that the first copy is already mutated in all the cells in the organism (Donehower et al. 2019). Despite the high cancer prevalence, relatively little is known about the early somatic evolutionary processes that predate and ultimately lead to LFS carcinogenesis, partly because LFS is rare and partly because most sequencing efforts profile cancers rather than normal tissue (Light et al. 2023; Boruah et al. 2024). Recent deep-sequencing studies capable of detecting cellular clones composed of only a few cells have revealed new insights into the mutational and selective forces operating across diverse healthy tissues in the general population, shedding light on how somatic variation arises and can later drive cancer development (Kakiuchi and Ogawa 2021; Salk et al. 2019; Calvet et al. 2025; Lawson et al. 2025). Similar efforts to quantify somatic evolutionary processes in individuals with LFS could lead to a better understanding of their risk of cancer development and inform diagnostic and monitoring strategies.

Deep sequencing of normal tissue in individuals with LFS offers a unique approach to understand how a single altered copy of *TP53* affects somatic evolution across tissues. Somatic *TP53* mutations are a common first hit across numerous cancer types in the general population (Martínez-Jiménez et al. 2020), and they may accelerate mutagenesis or clonal selection elsewhere in the genome. Because all cells in individuals with LFS carry a germline *TP53* variant, studying somatic evolution in these individuals allows a comprehensive examination of how second-hit mutations arise and expand. Further, when *TP53* pathogenic variants are present in the germline, carrier cells exist in all anatomical locations, providing an opportunity to understand how subsequent somatic evolutionary processes differ across tissue contexts and predispose LFS individuals to specific cancer types (McBride et al. 2014).

However, the detection of somatic mutations in normal tissue is challenging because those mutations are rare and escape detection by conventional sequencing methods. Duplex sequencing is an error correction sequencing method that enables ultradeep sequencing (>10,000×) and unprecedented resolution for mutation detection (Kennedy et al. 2014). In this Resource paper, we use duplex sequencing of multiple genomic regions to compare somatic evolution patterns in carriers and non-carriers of the *TP53* germline pathogenic variant c.542G>A (p.Arg181His or p.R181H hereafter). Among inherited *TP53* pathogenic variants, p.R181H is notable for its relatively high prevalence, partial loss-of-function phenotype, reduced cancer penetrance, and later cancer onset compared to other LFS-associated pathogenic variants (Timofeev and Stiewe 2021; Klimovich et al. 2022; de Andrade et al. 2024; Kratz et al. 2021). Despite attenuated effects, individuals with germline partial loss-of-function *TP53* variants such as p.R181H maintain substantially increased cancer risk compared to the general population, developing cancer at younger ages, especially breast cancer (Sun Zhang et al. 2025). The p.R181H variant has also been associated with cancers outside the main LFS spectrum, including prostate, colorectal, and thyroid (de Andrade et al. 2024; Freycon et al. 2024; Monti et al. 2007; Bougeard et al. 2015). One individual with LFS in our cohort developed esophageal cancer at age 34 and succumbed to metastatic disease. Multi-tissue collection at autopsy and blood samples procured from their family members and healthy donors provided a valuable opportunity to investigate mutagenesis and clonal expansions in the context of the *TP53* p.R181H germline variant.

## Results

### Ultra-deep sequencing of a family carrying p.R181H and healthy controls

The study included an individual with LFS who carried the germline *TP53* p.R181H pathogenic variant and died of metastatic esophageal carcinoma at age 34, as well as members of his family. Ultradeep DNA duplex sequencing of autopsy-collected tissues provided a unique opportunity to investigate how the germline p.R181H status influences somatic evolution and clonal dynamics in blood and solid tissues. Our cohort included the proband (LFS01), two family members carrying the same pathogenic variant and not known to have cancer (LFS02, LFS03), one relative without the pathogenic variant (REL01), and seven additional unrelated healthy controls of variable age between 25 and 76 ((CON01-CON07), **Figure 1A, Supplemental Table S1**).

**Figure 1.**
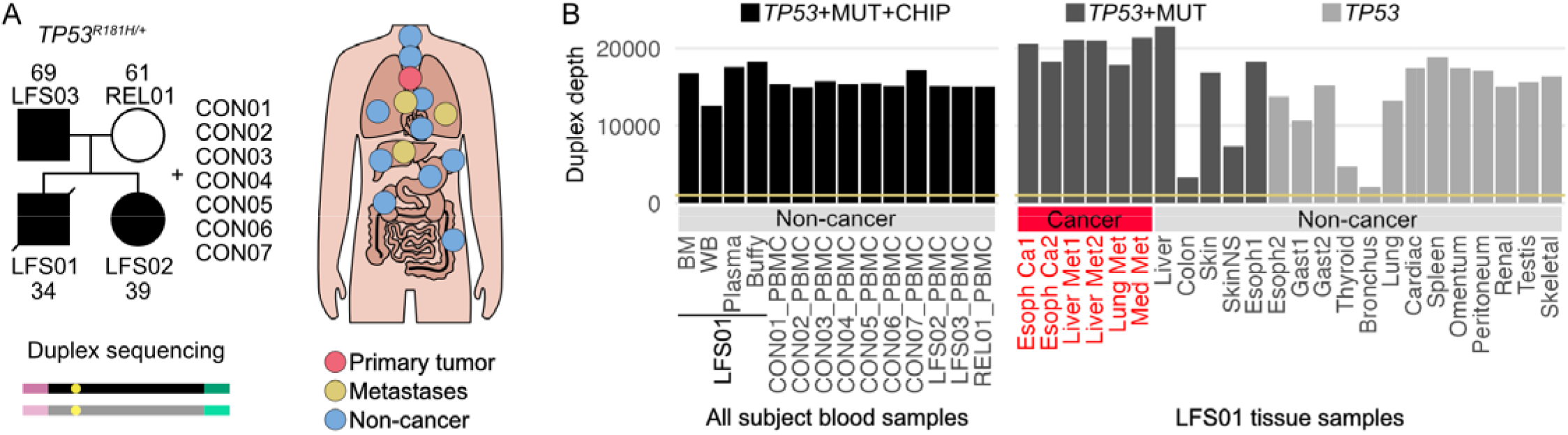
Ultra-deep sequencing of somatic mutations in subjects with and without Li-Fraumeni Syndrome. (A) The study included a Li-Fraumeni patient (LFS01, proband), his family (REL01, LFS01-02), and 7 unrelated controls (CON01-07). The pedigree indicates germline *TP53* p.R181H pathogenic variant carriers and ages at sampling. Blood from all individuals, along with primary esophageal (red), metastatic (yellow), and non-cancer (blue) tissues collected from the proband at autopsy were analyzed with ultra-deep duplex sequencing. This method uses double stranded molecular tags that enable mutation calling i both strands of DNA independently. *Tissues not shown in the diagram include skeletal muscle (right pectoralis), sun-exposed skin (left shin), non-sun-exposed skin (left buttock), omentum, peritoneum, and testis. (B) Duplex sequencing was performed with 3 different panel combinations depending on sample: *TP53* panel only, *TP53* panel with the mutagenesis panel (MUT), or *TP53* panel combined with both the MUT and the AML-29 Assay (CHIP) panel. Duplex depth (calculated as the mean number of duplex reads per position) is shown for each sample. For some tissues, two samples (labeled 1 or 2) were analyzed. Yellow line represents 1000x duplex depth sample threshold. BM: bone marrow; WB: whole blood; PBMC: peripheral blood mononuclear cells; Esoph: esophagus; Ca: cancer; Met: metastasis; Med: mediastinal; SkinN : non-sun-exposed skin; Gast: gastric tissue.

Hematopoietic samples were collected from all study participants: buffy coat, plasma, whole blood, and bone marrow from LFS01 and peripheral blood mononuclear cells (PBMCs) from all other subjects (**Figure 1B, Supplemental Table S2**). Low frequency mutations were detected on all hematopoietic samples using ultradeep duplex sequencing with two targeted panels: 1) TwinStrand’s AML-29 Assay (Oshima et al. 2024), a panel of 29 genes recurrently mutated in acute myeloid leukemia (AML) and clonal hematopoiesis of indeterminate potential (CHIP, 59 kb footprint, **Supplemental Table S3**), referred to as CHIP genes hereafter; and 2) Twinstrand’s Human Mutagenesis Assay (Axelsson et al. 2024), which contains 20 genomic regions selected to be an unbiased footprint of human genomic sequence contexts, avoiding repeats and homopolymers, and with no known role in human cancer (48 kb footprint). The mean duplex consensus sequencing depth (duplex depth) for hematopoietic samples was 15,683× (12,606× - 18,282×, **Figure 1B**), with a cumulative duplex depth coverage for all samples of 219,408×. Most targeted genes and regions showed consistently high depth across all samples (**Supplemental Fig. S1A-B, Supplemental Table S4**). The extremely high depth of sequencing with these target panels enabled us to characterize how the p.R181H variant impacts clonal evolution and mutagenesis in the blood. Unless otherwise specified, all reported sequencing depths and mutation frequencies refer to duplex consensus sequences.

We also conducted extensive multi-tissue sampling in LFS01 (**Figure 1A**). Two primary esophageal tumor samples and four metastatic samples across three metastatic sites (liver, lung, and mediastinal) were collected. In addition, an extensive range of 18 non-cancerous samples spanning 16 tissues were collected (**Supplemental Table S2**). Duplex sequencing of *TP53* was performed in all samples, and a subset of 11 tissues (all tumor and metastases plus five non-cancer samples: esophagus, liver, colon, sun-exposed skin, and non-sun-exposed skin) were sequenced with the mutagenesis panel. The mean duplex depth was 15,264×, with a cumulative duplex depth coverage of 361,302×. Four tissue samples had reduced coverage with depths between 2,118-7,376× (thyroid, bronchus, colon, and non-sun-exposed skin, **Figure 1B**, **Supplemental Table S4**), which was due to DNA loss during library preparation. Across all individuals, tissue types, and sequencing panels, we identified 1,992 somatic coding mutations in CHIP genes including *TP53* and 6,487 somatic mutations in the mutagenesis panel (**Supplemental Tables S5-S7**). Sequencing also confirmed that all tissues from affected individuals were heterozygous for *TP53* R181H.

### Germline p.R181H does not affect most CHIP clonal expansion but is associated with increased mutation frequency in the mutagenesis panel and reduced positive selection in *TP53*

We first examined if the germline p.R181H variant altered the number of mutations acquired in CHIP genes and the degree to which cells carrying those mutations could clonally expand. To do so, we analyzed duplex sequencing of 29 genes associated with CHIP and/or AML in hematopoietic samples, using buffy coat from LFS01 as the closest comparable sample to the PBMCs analyzed from all other subjects. The number of mutations in the coding regions of CHIP genes varied across individuals, with more mutations found in samples from older individuals and in the blood of LFS01 (**Figure 2A**). However, this excess of mutations could be due to multiple confounders, including higher sequencing depth and exposure to chemotherapy. LFS01 was exposed to multiple chemotherapy treatments including 5-fluorouracil, which is mutagenic, as well as other non-mutagenic agents (see Methods). CON07 was exposed to cyclophosphamide, vincristine, and doxorubicin, which are agents without significant SNV mutagenic effects but with potential to promote expansion of preexisting mutant clones (Mitchell et al. 2025; Uryu et al. 2025). To control for differences in sequencing depth, for each sample we calculated CHIP Mutation Frequency (MF) as the number of coding mutations identified in CHIP genes divided by the total number of nucleotides sequenced in the coding regions of those genes. CHIP MF significantly increased with age (**Figure 2B**). The association with age was also significant for mutations in *DNMT3A* and *TET2* (**Supplemental Fig. S2**), which are some of the most common CHIP genes in the population (Kessler et al. 2022). Multivariate analysis demonstrated that CHIP MF was associated with age and chemotherapy but not with the germline p.R181H variant **(Figure 2C, Supplemental Table S8**). To determine if the p.R181H variant affected not just the number of mutant clones but also their degree of clonal expansion, we quantified CHIP Mutation Burden (MB), which takes into consideration the number of mutations and the clone size. CHIP MB was calculated as the total number of duplex reads with coding mutations in any CHIP gene divided by the total number of coding nucleotides sequenced. CHIP MB was significantly associated with age in univariate and multivariate models but it was not associated with chemotherapy or the p.R181H variant (**Supplemental Fig. S3, Supplemental Table S8**).

**Figure 2.**
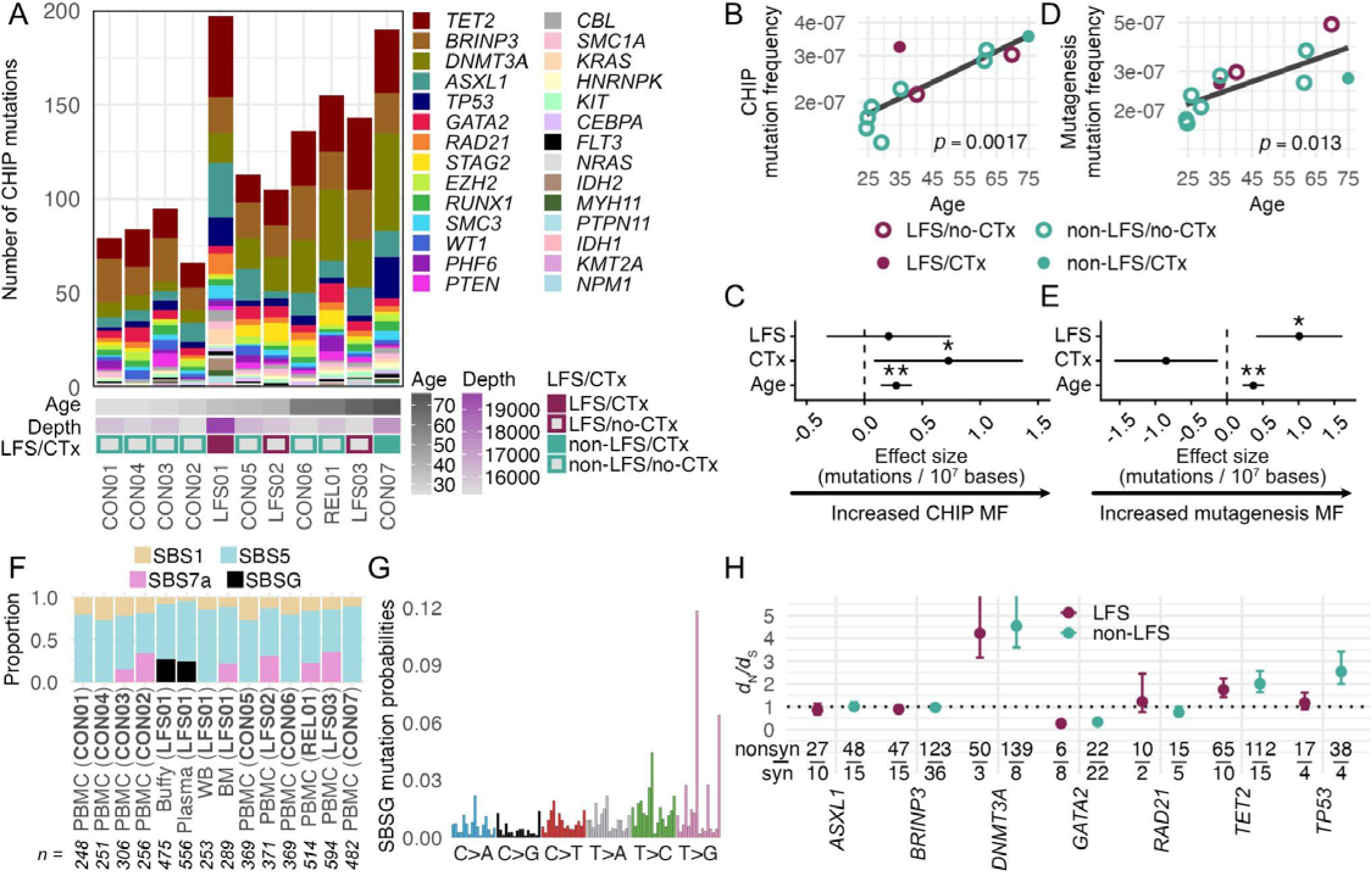
Ultra-deep characterization of CHIP mutation and selection in the blood of individuals with and without Li-Fraumeni Syndrome. (A) Number of coding mutations in CHIP/AML-associated genes by individual. Heatmap indicates age and average duplex sequencing depth across gene panel. Li-Fraumeni Syndrome (LFS) and chemotherapy history (CTx) are indicated by the color of the squares. (B,D) Linear regressions of coding CHIP mutation frequency (MF) (B) and mutagenesis panel MF (D) with age. Circles are colored by LFS status and filled circles indicate CTx history. Age-associated baseline regression is fit using all individuals and linear regression p-value is indicated. (C,E) Multiple regressions of coding CHIP MF (C) and mutagenesis panel MF (E) using age (scaled by decade), LFS, and CTx status as covariates. MF was scaled per 10,000,000 bases. Asterisks indicate significance value (*p<0.05, **p<0.01) and error bars indicate 95% confidence intervals. (F) Proportion of mutations associated with a mutational signature using mSigHDP by subject and tissue. The number of mutations within each sample is listed beneath each bar. (G) Mutational spectra of SBSG. The full 96 trinucleotide substitution spectra are shown, but only the six base substitution classes are labeled. (H) *d*_N_/*d*_S_ grouped by gene and germline *TP53* p.R181H status. Error bars indicate the interquartile range of *d*_N_/*d*_S_ values obtained through bootstrap resampling. Error bars extend beyond the plotted range for *DNMT3A*. Fraction indicates the number of observed non-synonymous (top) and synonymous (bottom) mutations used in calculation. Dotted line indicates absence of selection (*d*_N_/*d*_S_ =1).

In addition to CHIP genes, hematopoietic samples were also sequenced with a mutagenesis panel covering twenty 2.4 kb targets avoiding genomic regions under strong selection to serve as a genome-representative cross section of mutational contexts (Axelsson et al. 2024). Somatic mutations identified in those regions were used to calculate a MF (see Methods) called “mutagenesis MF” to distinguish it from CHIP MF. As expected, mutagenesis MF was significantly associated with age in univariate and multivariate models (**Figure 2D-E, Supplemental Table S8**). Interestingly, it was also associated with the germline p.R181H variant but not with chemotherapy, indicating that the chemotherapy treatments of the patients in the study did not have substantial effects as mutagens. Overall, these results indicate that the germline p.R181H variant does not globally elevate CHIP mutation frequency or burden but it is associated with higher mutation frequency in unselected genomic regions (**Figure 2E**).

Given that the p.R181H variant status was associated with elevated mutagenesis MF, we considered that the presence or intensity of active mutational signatures may differ based on carrier status. We performed *de novo* single base signature extraction using Mutational Signature Hierarchical Dirichlet Process (mSigHDP, (Liu et al. 2023)), a Bayesian model employed for de novo extraction of mutational signatures, on the mutations identified with the CHIP and mutagenesis panels (**Figure 2F**). Additional hematopoietic samples were analyzed for mutational signatures including plasma, whole blood, and bone marrow from LFS01. Extracted signatures resembled COSMIC signatures SBS1, SBS5, SBS7a and one signature (reported here as SBSG, **Figure 2G**) previously linked to 5-fluorouracil chemotherapeutic exposure (Mitchell et al. 2025). Clocklike signatures SBS1 and SBS5 were detected in all samples, with similar relative contribution in p.R181H carriers vs non-carriers (mean SBS1 and SBS5 proportion in carriers are 10.5% and 63.8% vs 19.5% and 71.6% in non-carriers, respectively). SBS7a was discovered in five individuals. SBS7a is caused by exposure to ultraviolet light and potentially reflects skin resident T cells recirculating into the blood (Machado et al. 2022). All but one sample containing SBS7a also had CC>TT dinucleotide variants characteristic of UV exposure (**Supplemental Fig. S4**). Two samples from LFS01 (buffy coat and plasma) showed evidence of chemotherapeutic signature SBSG, as expected due to 5-fluorouracil exposure for cancer treatment. SBSG was not detected in whole blood and bone marrow possibly due to limited resolution with lower mutation counts. No other p.R181H carrier-specific mutational signatures were detected across the affected individuals. However, we further tested for differences between the underlying mutational profiles found in p.R181H carriers and non-p.R181H carriers using a novel signature-agnostic metric called “aggregate mutation spectrum distance” permutation method (AMSD), which determines whether mutational exposures differ between groups (see Methods and (Hart et al. 2026)). The AMSD test found a small but significant difference between signatures based on carrier status (cosine similarity of 0.919, 0.032), potentially driven by the presence of SBSG in LFS01 or differences in SBS7a contribution. We therefore conclude that while the p.R181H variant may be associated with minor alterations in mutational processes, no single mutational signature clearly explains the difference.

Next, we examined if the p.R181H variant might affect positive selection in specific CHIP-associated genes, even if it did not appear to associate significantly with clonal expansions across the panel as a whole. For genes with at least 10 mutations in p.R181H carriers and non-carriers, we computed *d*_N_/*d*_S_ in both groups to look for evidence of carrier status-specific positive selection. *d*_N_/*d*_S_ revealed positive selection in *DNMT3A* and *TET2* and negative selection in *GATA2*, but these findings were comparable between p.R181H carriers and non-carriers. Interestingly, the only gene with non-overlapping bootstrapped interquartile ranges between the two groups was *TP53* (**Figure 2H**), which showed no selection in p.R181H carriers but positive selection in non-carriers. Pathogenicity analyses did not show significant differences in pathogenicity scores or percentage of pathogenic mutations for each gene, including *TP53*, between p.R181H carriers and non-carriers (**Supplemental Fig. S5**). The gene with the most pathogenic mutations was *DNMT3A* and the gene with the least pathogenic mutations was *GATA2*, supporting the *d*_N_/*d*_S_-based evidence of strong, but p.R181H independent, positive and negative selection in these genes, respectively (**Figure 2H**). While positive selection for *DNMT3A* is well-recognized (Kessler et al. 2022), the negative selection of *GATA2* has not been observed in population-based somatic sequencing and highlights a striking divergence in selective pressures across genes operating within healthy tissues. Nevertheless, the selection dynamics on CHIP genes in the blood is mostly unchanged by germline p.R181H status with the exception of *TP53* mutations, which are not selected in p.R181H carriers but are positively selected in non-carriers.

### Chemotherapy increases *TP53* clonal expansions in the blood of a germline p.R181H carrier and non-carrier

Given the central role of *TP53* in LFS and the observed differences in *TP53 d*_N_/*d*_S_ between LFS and non-LFS groups, we further characterized the individual somatic *TP53* mutations identified in blood within this cohort. Each individual carried between 2 and 20 *TP53* coding mutations in blood, with the highest counts observed in the two individuals with previous chemotherapy exposure (LFS01 and CON07, **Figure 3A**). Most mutations appeared to be pathogenic based on mutation type and the AlphaMissense algorithm (see Methods, **Figure 3A**) and they clustered in the DNA binding domain of the protein (75% compared to 46% expected, binomial test, 10 **Figure 3B-C**). The observed missense mutations, especially those seen on more than one duplex read, were enriched for higher pathogenicity scores compared to the distribution of scores for all possible *TP53* missense mutations (**Figure 3D**, 0.0012). These results suggest that the *TP53* mutations observed in the blood mostly correspond to positively selected pathogenic somatic mutations.

**Figure 3.**
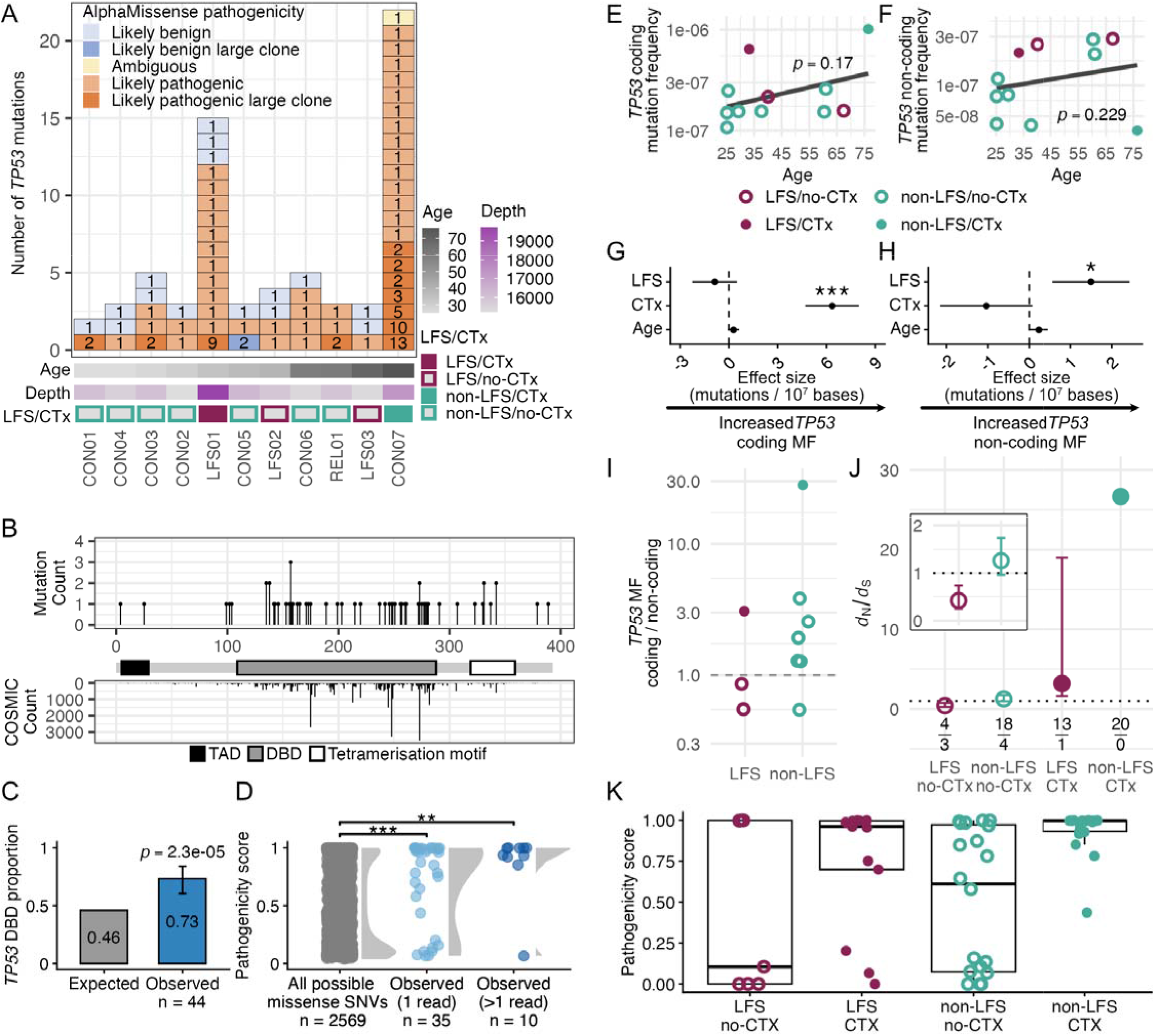
Ultra-deep characterization of *TP53* mutation and selection in the blood of individuals with and without Li-Fraumeni Syndrome. (A) Number, size, and pathogenicity of coding *TP53* mutations by individual. Each box represents a unique *TP53* mutation within an individual. Box color indicates pathogenicity based on mutation type and AlphaMissense scores (see Methods). The number inside of each box represents the number of duplex reads supporting the mutation. Mutations seen in more than 1 duplex read are considered large clones. Heatmap indicates age and average duplex sequencing depth of *TP53*. Li-Fraumeni Syndrome (LFS) and chemotherapy history (CTx) are indicated by the color of the squares. (B) Single nucleotide variant (SNV) counts within the *TP53* coding region across all individ als (top) and mutation counts in the COSMIC database (bottom) by codon position. Major functional regions shown: topologically associating domain (TAD), DNA-binding domain (DBD), and tetramerization motif. (C) Expected and observed SNV proportion within the DNA-binding domain of *TP53*. Expected proportion corresponds to the DNA-binding domain’s base pair proportion of the total *TP53* coding region. Binomial test p-value and error bars representing 95% confidence interval are shown. (D) AlphaMissense pathogenicity scores of all possible *TP53* missense mutations, missense mutations observed in one duplex read, and missense mutations observed in >1 duplex read. p-values correspond to Mann-Whitney tests with asterisks indicating significance value (*p<0.05, **p<0.01, ***p<0.0001) (E,F) Linear regressions of coding *TP53* mutation frequency (MF) (E) and non-coding *TP53* MF (F) with age. Circles are colored by LFS status and filled circles indicate CTx history. Age-associated baseline regression is fit using all individuals and linear regression p-value is indicated. (G,H) Multiple regressions of coding *TP53* MF (G) and non-coding *TP53* MF (H) using age (scaled by decades), LFS, and CTx status as covariates. For MF calculations, sequencing depth was scaled per 10,000,000 bases. Asterisks indicate significance value (*p<0.05, **p<0.01, ***p<0.0001) and error bars indicate 95% confidence intervals. (I) Ratio of *TP53* coding and non-coding MF by LFS status. (J) *TP53 d*_N_/*d*_S_ grouped by germline p.R181H status and CTx status. Error bars indicate the interquartile range of *d*_N_/*d*_S_ values obtained through bootstrap resampling. Fraction indicates the number of observed non-synonymous (top) and synonymous (bottom) mutations used in calculation. Inset shows a magnified view of the no-CTx groups only. (K) AlphaMissense pathogenicity scores of observed *TP53* SNVs by germline p.R181H status and CTx status.

Given that observed *TP53* coding mutations appear to be positively selected, we next calculated *TP53* coding and non-coding MF in blood samples (the latter counting only non-coding mutations captured by the *TP53* panel, **Supplemental Table S9**) to better resolve the contribution of selection and baseline mutagenesis respectively. Similar to the CHIP-panel results in aggregate, *TP53* coding and non-coding MF increased with age, although not significantly (**Figure 3E-F**). In multivariate models, *TP53* coding MF was only significantly associated with chemotherapy whereas *TP53* non-coding MF was associated with p.R181H status (**Figure 3G-H, Supplemental Table S8**), mirroring the associations observed in the CHIP panel more broadly. To enable a better understanding of the relationship between *TP53* coding and non-coding MF based on LFS status, we calculated the ratio of *TP53* coding to non-coding MF for each individual (**Figure 3I**). We observed that chemotherapy increases the ratio regardless of LFS status, concordant with chemotherapy promoting the expansion of *TP53* mutant clones. Additionally, we observed that most individuals without LFS and no chemotherapy had a ratio >1 whereas the two individuals with LFS and no chemotherapy had a ratio <1. These results agree with the prior observation of lack of positive selection of *TP53* mutations in the blood of LFS individuals (**Figure 2H**) and indicate that, to properly compare *d*_N_/*d*_S_ between groups, the individuals with chemotherapy should be considered separately. Thus, we recalculated the *d*_N_/*d*_S_ for *TP53* mutations in blood by separating LFS and chemotherapy status (**Figure 3J**). We observed that chemotherapy greatly increased *d*_N_/*d*_S_ across both LFS and non-LFS relative to their respective no chemotherapy groups. This supports chemotherapy as a promoter of the expansion of clones with *TP53* pathogenic mutations. Without the confounding of chemotherapy, we observed *d*_N_/*d*_S_ < 1 in LFS individuals indicating negative selection (**Figure 3J**). Consistent with the *d*_N_/*d*_S_ result, pathogenicity scores demonstrated high levels for individuals with chemotherapy and lower levels for individuals without chemotherapy with the lowest proportion of *TP53* pathogenic mutations found in LFS individuals without chemotherapy (**Figure 3K**).

In summary, the two LFS individuals without chemotherapy (LFS02 and LFS03) had high levels of non-coding mutations and no comparable increase of *TP53* coding mutations. Only 4/7 (57%) of the *TP53* coding mutations found were non-synonymous compared with 18/22 (82%) in non-LFS patients. While the difference was not statistically significant, it explains the lower *d*_N_/*d*_S_ values observed in these individuals. Overall, these results suggest that the p.R181H germline variant might be associated with an increase in mutagenesis but a reduction in the positive selection of *TP53* pathogenic clones. Unfortunately, the strong selective action of chemotherapy precludes confirmation of this finding in the blood of the treated patient.

### Mutation frequency and signatures vary across tissue contexts

A unique feature of this study was the availability of both normal and cancerous tissue from 22 different anatomical locations (**Figure 1A-B**), which enabled us to explore *TP53* somatic evolution across different tissues in the context of a germline p.R181H pathogenic variant. We discovered considerable variation in the number of somatic *TP53* mutations in different tissues (**Figure 4A**). The majority of these mutations were missense, many of which were classified as pathogenic by AlphaMissense (**Figure 4A**). Sun-exposed skin contained more than twice as many *TP53* mutations as the next most mutated sample. Many of these mutations were CC>TT dinucleotide variants, likely related to UV-induced mutagenesis (**Figure 4B**). Four samples had disproportionately low sequencing depth compared to the rest (**Figure 4A**), which likely was responsible for their low mutation count. These samples were discarded for mutation frequency analyses. For the rest of the samples, we calculated *TP53* coding and non-coding MF (**Figure 4C, Supplemental Table S10**). With the exception of sun-exposed skin, samples had relatively low and homogeneous levels of non-coding MF (mean: 1.37 10). *TP53* coding MF was low and comparable to non-coding MF in bone marrow, peritoneum, testis, kidney and some cancerous samples. In contrast, *TP53* coding MF was elevated in sun-exposed skin, peripheral blood samples (whole blood, buffy coat and plasma), and normal esophagus, which was the tissue of origin of the cancer. Interestingly, the patient had chronic gastritis and *TP53* coding MF was also elevated in the two normal gastric samples. The primary and metastatic cancer samples did not have higher *TP53* coding MF compared to healthy tissues. All cancer and metastatic samples, however, had loss of heterozygosity (LOH) of the wild-type *TP53* allele as reflected by variant allele frequencies >0.5 for the germline p.R181H variant (**Supplemental Fig. S6**). These results indicate that the loss of the wild-type allele likely drove tumorigenesis in this individual. LOH-driven carcinogenesis may also explain the lower coding MF of the primary tumor compared to non-cancer esophagus as biallelic *TP53* inactivation may eliminate the selective advantage for additional *TP53* pathogenic mutations. Several non-cancer tissue samples (lung, bronchi, esophagus, omentum, and gastric tissue) also showed variant allele frequencies >0.5 for p.R181H. While these values could represent early LOH expansion, these biopsies were in proximal location to the primary and metastatic sites and pathological findings were consistent with cancer dissemination (micrometastases) in some of the samples, making this the most likely explanation for increased variant allele frequency (**Supplemental Fig. S6**).

**Figure 4.**
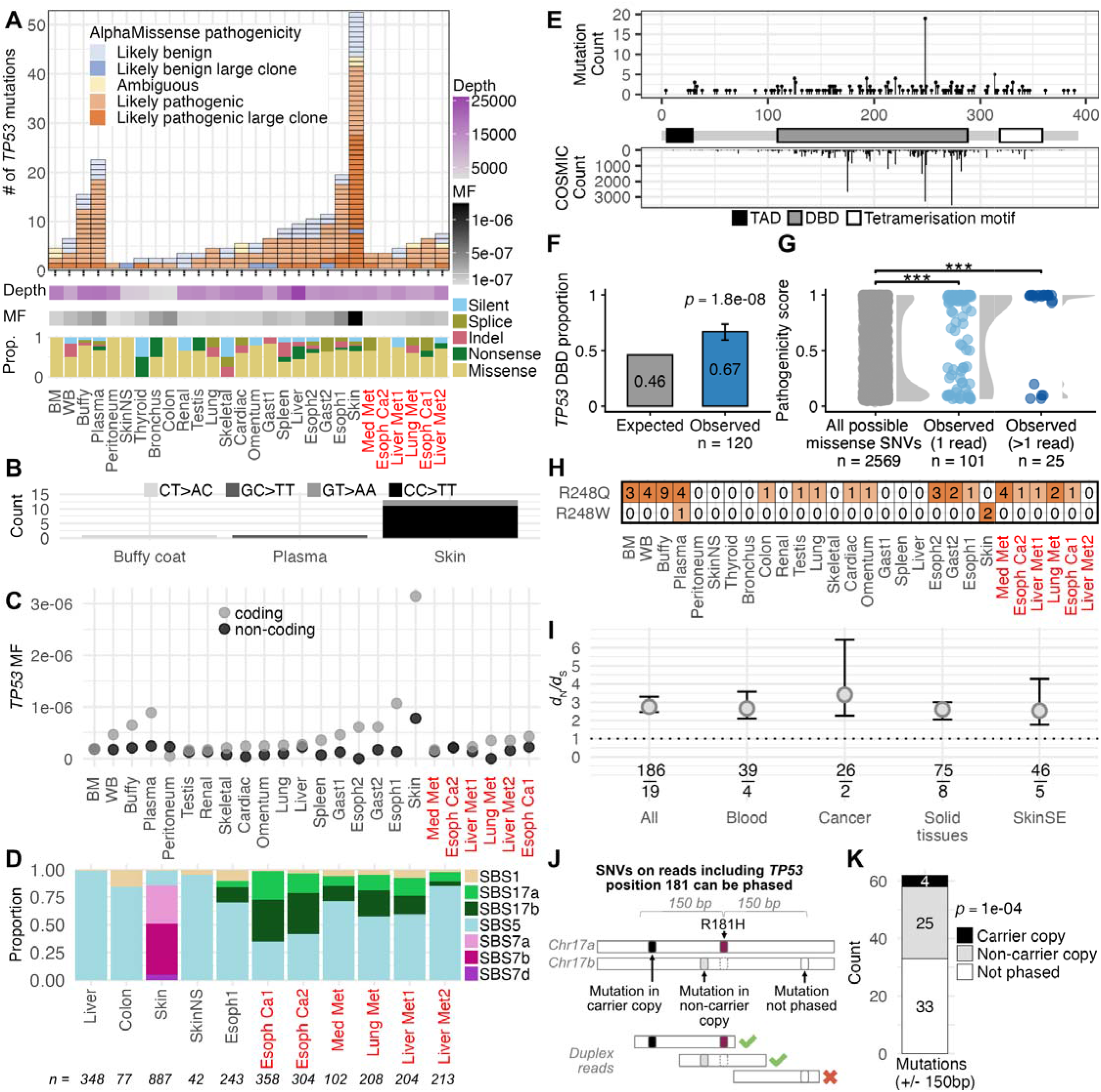
Multi-tissue ultra-deep characterization of TP53 mutation and selection in an individual with Li-Fraumeni Syndrome and esophageal adenocarcinoma (LFS01). (A) Number, size, and pathogenicity of coding *TP53* mutations by tissue in LFS01. Each box represents a unique *TP53* mutation within a tissue. Box color indicates pathogenicity based on mutation type and AlphaMissense score (see Methods). Mutations seen in more than 1 duplex read are considered large clones. Heatmaps indicate duplex depth (purple) and mutation frequency (MF, black). Bar plot represents the proportion of *TP53* mutation type by tissue. Cancer samples indicated in red text. (B) *TP53* dinucleotide variant counts by tissue in which they were observed. (C) *TP53* coding and non-coding MF by tissue. (D) Proportion of mutations associated with a mutational signature using mSigHDP. Only samples sequenced with *TP53*+MUT panel were included. Cancer samples indicated in red text. The number of mutations within each sample is listed beneath each bar. (E) Single nucleotide variant (SNV) counts within the *TP53* coding region across all tissues in LFS01 (top) and mutation counts in the COSMIC database (bottom) by codon position. Major functional regions shown: topologically associating domain (TAD), DNA-binding domain (DBD), and tetramerization motif. (F) Expected and observed SNV proportion within the DNA-binding domain of *TP53*. Expected proportion corresponds to the DNA-binding domain’s base pair proportion of the total *TP53* coding region. Binomial test p-value and error bars representing 95% confidence interval are shown. (G) AlphaMissense pathogenicity scores of all possible *TP53* SNVs, SNVs observed in one duplex read, and SNVs observed in >1 duplex read. p-values correspond to Mann-Whitney tests with asterisks indicating significance value (*p<0.05, **p<0.01, ***p<0.0001) (H) Size and tissue location of *TP53* mutations by amino acid substitution at codon position 248. The number inside of each box represents the number of duplex reads associated with that given mutation. Cancer samples indicated in red text. (I) *TP53 d*_N_/*d*_S_ grouped by tissue categories, with all tissues aggregated within All. Error bars indicate the interquartile range of *d*_N_/*d*_S_ values obtained through bootstrap resampling. Fraction indicates the number of observed non-synonymous (top) and synonymous (bottom) mutations used in calculation. (J) Phasing analysis schematic. The chromosome copy carrying p.R181H and non-carrier copy may acquire new somatic mutations. When a somatic mutation is observed on the same duplex read as the germline mutation site, the allelic phase relative to p.R181H can be determined. (K) Number of somatic *TP53* mutations phased on the carrier copy, non-carrier copy, or not capable of being phased across all three LFS individuals (LFS01, LFS02, LFS03). Binomial test significance between alleles indicated with p-value.

Mutational signature extraction and decomposition across a subset of the sampled tissues further supported variation in active mutational processes in different tissue contexts (**Figure 4D**). All tissues contained clocklike mutational signatures SBS1 and SBS5, with SBS5 being more prevalent than SBS1 in all tissues. As expected, mutations in sun-exposed skin were dominated by UV-associated signatures SBS7a/b/d. Both primary esophageal cancer samples, four metastatic cancer samples and one healthy esophageal sample showed contributions from SBS17a/b, which have previously been linked to esophageal carcinoma and premalignant Barrett’s esophagus (Abbas et al. 2023). SBS17b has also been associated with 5-fluorouracil treatment (Pich et al. 2019), making it difficult to unambiguously disentangle the observed drivers of mutagenesis. The high resemblance in mutational signatures between replicate samples (two esophageal cancer samples and two liver metastases) demonstrates the robustness of our approach for high resolution mutation detection and signature extraction and decomposition.

### *TP53* R248Q/W is a mutational hotspot recurrently found in several tissues

A powerful attribute of examining multiple tissues from a single individual is the ability to characterize evolutionary processes of particular genetic alterations occurring independently in matched genetic backgrounds, potentially indicating their selective importance. We therefore examined if the observed mutations preferentially clustered in particular regions of *TP53*. The majority of discovered SNVs were observed in the DNA-binding domain, similar to *TP53* mutations in COSMIC as a whole (**Figure 4E**), our findings in blood (**Figure 3B**), and in prior studies of somatic *TP53* mutations in normal tissue (Salk et al. 2019; Rios-Doria et al. 2025; Matas et al. 2022). The enrichment of *TP53* mutations in the DNA-binding domain was statistically significant (**Figure 4F**) indicating positive selection of functionally disruptive mutations as a second hit in the context of a germline p.R181H background. In addition, observed mutations were enriched for higher AlphaMissense scores than expected compared to the distribution of scores for all possible *TP53* missense mutations, especially if the mutations were supported by more than one duplex read, indicating the expansion of pathogenic clones (**Figure 4G**,.). Strikingly, we found that 18/28 tissue samples contained a mutation at codon 248, the second most widely-mutated *TP53* hotspot in the COSMIC database (**Figure 4E,H)**. We observed two different amino acid changes (p.R248Q and p.R248W), both of which are inactivating and dominant negative (Balasundaram and Doss 2022; Klemm et al. 2025; Boettcher et al. 2019). The p.R248Q was observed more frequently than the p.R248W across tissue contexts, although the plasma sample contained both simultaneously (**Figure 4H**). While these mutations remained at low variant allele frequencies that would be undetectable by standard sequencing methods, 9/19 were observed in >1 duplex read, indicating some degree of clonal expansion. While it is possible for leukocyte clones to infiltrate the tissues and be sampled and detected by ultra-deep sequencing, it is unlikely for a clone of such low variant allele frequency (the largest clone in buffy coat has a frequency of 0.0005) to repeatedly be sampled across different tissues. In addition, we found little evidence that biological or technical contamination was responsible for this widespread mutational sharing, as most other mutations in the blood were not found across tissues (**Supplemental Fig. S7**). The second most frequent mutation was p.S314F, which was found only in four tissues (**Supplemental Fig. S8**). These results broadly suggest convergent evolution favoring disruption of the arginine at position 248 across diverse tissue contexts in the background of a germline p.R181H variant.

### Allelic specificity but not tissue-context accounts for differences in selection pressures in *TP53*

We next examined if *TP53* mutations were subject to different selective pressures across tissue contexts. To ensure sufficiently large sample sizes for robust computation of *d*_N_/*d*_S_, we aggregated samples into the larger anatomical categorizations of blood, solid tissues, sun-exposed skin, and cancer (**Supplemental Table S11**). Consistent with our blood findings for LFS01 (**Figure 3J**), we found significant elevation of *d*_N_/*d*_S_ across multiple tissue contexts, suggesting positive selection for secondary *TP53* mutations (**Figure 4I**). Given the negative selection for *TP53* mutations observed in the blood of p.R181H carriers without chemotherapy, it is likely the positive selection observed in these samples to be a consequence of chemotherapy exposure. The magnitude of *d*_N_/*d*_S_ was similar across tissue groupings, although these groupings may mask more localized tissue effects.

Finally, using the large sample size of *TP53* mutations observed across tissues in all LFS individuals within our cohort, we asked if the chromosomal context of *TP53* mutations affected their ability to be selected. Specifically, we hypothesized that there might be greater positive selection for *TP53* mutations occurring on the chromosomal copy not already carrying p.R181H, as opposed to the partially inactivated p.R181H carrier copy (**Figure 4J**). The relatively short length of duplex reads limited the number of somatic mutations that could be confidently phased to 29. Of these mutations, significantly more (25/29; binomial test, 0.0001) occurred on the chromosomal copy not already carrying p.R181H, indicating increased selection for second hit mutations affecting the non-carrier *TP53* copy (**Figure 4K**). This significant allelic imbalance supports ongoing selection for second-hit *TP53* inactivating mutations in cells with p.R181H.

## Discussion

Ultra-deep sequencing of normal tissues has revealed that somatic evolution is widespread (Kakiuchi and Ogawa 2021; Salk et al. 2019; Matas et al. 2022). Measuring mutational and selective processes across tissues, sexes, environmental exposures and populations is providing new understanding of the mechanisms driving cancer formation (Calvet et al. 2025; Lawson et al. 2025; Yokoyama et al. 2025). Here, we apply this ultra-deep sequencing approach to investigate somatic evolution in a family of individuals with LFS caused by the germline *TP53* pathogenic variant p.R181H, one of the most prevalent mutations causing cancer predisposition (de Andrade et al. 2024; Moses et al. 2025; Sun Zhang et al. 2025). We find that the germline p.R181H variant is associated with increased mutational frequency in the mutagenesis panel but not in the coding regions of CHIP and/or AML-related genes. A stark exception, however, is the study proband, who had a large number of mutations in CHIP panel genes likely related to chemotherapy exposure, which precluded disambiguation of the effect of the germline mutation vs the cancer treatment. Despite that limitation, we observed an interesting pattern of second hit mutations in the proband with widespread pathogenic substitutions in p.R248 across tissues. We also confirm expected patterns of somatic evolution in p.R181H carriers and non-carriers that suggest our analyses are robust, including the finding of chemotherapy mutational signatures in blood, UV light signatures in sun-exposed skin, and increased mutation frequency with age. In addition, despite a relatively young cohort, ultra-deep sequencing enabled the identification of abundant low frequency mutations in *DNMT3A* and *TET2*, the most common CHIP genes (Genovese et al. 2014; Marnell et al. 2021), which also showed strong evidence of positive selection. In contrast, we report the novel observation of strong negative selection in *GATA2*, which is consistent with its essential role in hematopoietic stem cell survival (Menendez-Gonzalez et al. 2019) and implies that even low-frequency deleterious variants are likely purged before clonal expansion can occur. Together, these findings underscore that the mutational landscape in healthy individuals already reflects lineage-defining constraints that may foreshadow later MDS/AML evolution, and that this landscape does not appear to be perturbed by the germline *TP53* p.R181H variant.

The increased frequency of mutations in the mutagenesis panel for LFS carriers indicates that p.R181H may affect mutagenesis. The primary phenotypic alterations of R181H mutants are their inability to transactivate p53 target genes and initiate cell cycle arrest, although these effects may be lessened in a heterozygous state (Moses et al. 2025). A potential result of this could be the accumulation of mutations in cells that would otherwise not proceed through division, although it is unclear why this would lead to an observed increase in the frequency of mutations in the mutagenesis panel, but not the CHIP panel. One possibility is that increased mutagenesis affects both panels, but the patterns in coding regions are obscured by the dynamics of selection. Importantly, the germline p.R181H mutation did not affect the selection of blood CHIP clones except for *TP53,* which had reduced selection in germline carriers. However, this finding is difficult to completely disambiguate from chemotherapy exposure in a subset of LFS carrier samples, so sequencing additional carriers without exposure (in addition to different tissue samples and genes) would be key to further explore these findings.

We found divergent strengths of positive selection on *TP53* mutations between the proband LFS01 and their relatives LFS02 and LFS03, all p.R181H carriers. The relatives showed reduced positive selection in *TP53*, as evidenced by a deficit of coding mutations, *d*_N_/*d*_S_ < 1, and lower predicted mutational pathogenicity compared to LFS01 and the non-p.R181H carriers profiled. In contrast, LFS01 showed abundant evidence for positive selection in *TP53* both in the blood and across tissues. A likely driver of these divergent dynamics is that LFS01 is the only relative to have received chemotherapy, which has been extensively shown to be associated with *TP53* clonal expansions (Takahashi et al. 2024; Bolton et al. 2020; Coombs et al. 2017; Nead et al. 2024). Finding reduced positive selection for *TP53* mutations among untreated individuals carrying the p.R181H variant may point to the importance of LOH rather than secondary SNVs to inactivate the second copy of *TP53*. It has been reported that loss of heterozygosity of *TP53* is an early and near-ubiquitous event in LFS-related cancers (Light et al. 2023) and LOH was indeed the second hit in the tumor of the study proband. While LOH might be the critical second hit in the progression to cancer, we did find some evidence that SNV-based *TP53* mutations are important in p.R181H carriers. We found widespread p.R248Q/W mutations across many tissue contexts in LFS01 and demonstrate that most SNVs occur in the alternate allele carrying the WT R181 amino acid. These results suggest that second hit *TP53* mutations do initiate clonal expansions but to what extent this is restricted to the selective pressure of chemotherapy is unknown.

Profiling cancerous and healthy tissue samples from distinct anatomical locations revealed some expected differences in the mutational and selective patterns of different organs in a p.R181H donor. Specifically, we observed greater number of CC>TT dinucleotide variants associated with UV-induced mutagenesis in the skin, and cancer-specific mutational signatures in tumor samples. We also observed that the biopsies with the highest levels of *TP53* mutation frequency, aside from sun-exposed skin, were those procured from the esophagus and the stomach. Interestingly, the patient had chronic acid reflux, which is a risk factor for esophageal cancer and chronic gastritis. Both conditions are associated with a carcinogenic process driven by *TP53* mutations, although in the esophagus, this typically involves Barrett’s metaplasia (Redston et al. 2022; Stachler et al. 2018) and in the stomach it is often related to *Helicobacter pylori* infection (Shimizu et al. 2014; Morgan et al. 2003). However, pathological assessment of the tissues did not identify Barrett’s esophagus or *H. pylori* infection in this patient. Nevertheless two independent biopsies of these two tissues showed high levels of *TP53* mutation supporting the link between chronic tissue damage and *TP53* clonal expansion. We could speculate that the germline p.R181H variant might have contributed to a faster carcinogenic process in the esophagus in the context of preexisting epithelial damage, but functional studies would be needed to test this hypothesis. Regarding selection, we did not observe differences across tissues using *d*_N_/*d*_S_ calculations. However, these calculations were performed by grouping mutations into tissue categories due to sample size considerations, which might have hidden associations present only in specific anatomical locations or cell types. In addition, these associations might also be overshadowed by the selective effect of chemotherapy. Access to normal tissue of LFS subjects without chemotherapy and sampling of multiple biopsies will be key to solve these questions in future studies.

Examining multiple tissues from a single individual also revealed widespread parallelism in the clonal evolutionary processes, especially for *TP53* mutations p.R248Q/W which were found in >60% of LFS01 tissues. These mutations are dominant negative, associated with increased cellular proliferation and reduced apoptotic response (Balasundaram and Doss 2022; Klemm et al. 2025; Boettcher et al. 2019; Olivier et al. 2010), and are widely mutated in human cancers. Given their prevalence in cancer, it may be unsurprising to observe their simultaneous expansions in multiple normal tissues. However, we did not see similar expansions at other putatively dominant negative *TP53* hotspots (i.e., R175H, Y220C, M237I, R273H, R282W), potentially pointing to a distinct interaction between p.R181H and p.R248Q/W. While this finding could be the result of technical or biological cross-contamination across samples, both possibilities seem unlikely given SNP quality control, the lack of other shared mutations across tissues, and the low variant allele frequency of p.R248Q/W in all samples. A possible explanation could be repeated mutations and expansions of p.R248Q/W across tissue contexts. Alternatively, two or more p.R248Q/W mutant leukocyte clones could be selectively infiltrating different tissues, an intriguing possibility that deserves further investigation. It would also be of interest to see whether similar preferences for second hits are observed for other *TP53* germline pathogenic variants.

The study had some limitations. First, while we had control samples for blood, we did not have control samples for the normal tissue, and available studies with comparable sequencing depths in concordant tissues are non-existent. Second, we had multiple tissues only from one individual and the individual was treated with chemotherapy, confounding potential selective dynamics in a p.R181H background. Inclusion of a chemotherapy exposed control (CON07) was useful to partially delineate the selective effects in blood, but further studies would benefit from sampling normal tissue in LFS patients prior to chemotherapy. Blood, buccal swabs, urine, and biopsies from colonoscopies are all feasible procedures for normal tissue collection. Lastly, tissues were only sequenced for *TP53* and a subset for the mutagenesis panel. We note that while this footprint is more limited than other studies of somatic evolution (Lawson et al. 2025), our sequencing depth is much greater (15,000x versus <1000x). This enables the discovery of rarer genetic variation within this young cohort that reveals new biological results, such as negative selection on *GATA2* and repeated observation of p.R248Q/W across tissue contexts. Now that sequencing is more affordable, extended panels are likely to provide more comprehensive information. Additionally, we only profiled one LFS pathogenic germline variant. While this is one of the more common germline *TP53* pathogenic variants, it also is unusual in its reduced penetrance, atypical cancer profile, and later age onset of cancer (Timofeev and Stiewe 2021; Klimovich et al. 2022; de Andrade et al. 2024; Kratz et al. 2021). Profiling somatic evolution in people with LFS with different germline *TP53* pathogenic variants will be necessary to understand the generalizability of our findings related to the increased non-coding mutagenesis, the lessening of *TP53* mutation selection, and the concordant second hits in p.R248. Other germline variants with higher penetrance might lead to different phenotypes that could be exploited to personalize surveillance and cancer prevention.

In summary, we demonstrate how ultra-deep sequencing can reveal the evolutionary processes operating in healthy and cancerous tissues of individuals carrying the germline *TP53* p.R181H variant. This study paves the way for other analyses in LFS patients and other cancer predisposition syndromes. We discovered distinct features of mutagenesis and selection in LFS tissues, which reflect underlying biological processes that are incompletely understood and deserve further investigation.

## Materials and Methods

### Sample collection

The study includes samples collected at autopsy from a male patient known to have LFS, who died from metastatic esophageal cancer at the age of 34 (23 normal tissues, 6 tumor tissues) (**Figure 1, Supplemental Tables S1 and S2**). The study also includes blood samples (PBMCs) collected from three family members and from 7 unrelated control donors (**Figure 1, Supplemental Tables S1 and S2**). Samples from the LFS-affected patient were collected through the willed body program at UW. The collection and analysis of samples from the patient, family members and donors was approved under IRB study number STUDY00020611.

Anatomic samples from the LFS-affected patient were collected at autopsy and immediately frozen at -80°C with an estimated post-mortem interval of 22h. The patient had a smoking history, chronic acid reflux, and chronic gastritis. Gastric tissue immunohistochemistry was negative for *Helicobacter pylori* and there was no evidence of goblet cell metaplasia or Barrett’s esophagus. The main tumor was moderately differentiated (grade II of III) adenocarcinoma in esophageal/gastric junction in a background of glandular dysplasia and ulceration. There was a satellite nodule in the cardia of the stomach 10 mm away from the lower border of the main cancer. The nodule had high-grade glandular dysplasia which is a precancerous lesion to adenocarcinoma. The patient had received chemotherapy consisting of folinic acid/5-fluorouracil/oxaliplatin (FOLFOX) and trastuzumab. As disease progressed, he was treated with a regimen consisting of ramucirumab/docetaxel/zoledronic acid and received palliative radiotherapy for metastatic bone lesions. None of the other subjects in the study had cancer except for the 76-year-old control donor, who had a history of non-Hodgkin lymphoma and was treated with two cycles of cyclophosphamide/doxorubicin/vincristine/prednisone (CHOP).

Frozen solid tissue samples were manually macrodissected using tissue punchers for a target size of 3mm^3^ or ∼ 25 mg of tissue. Single use tools were used to avoid cross-contamination between samples. Tissue DNA was extracted using the DNeasy Blood and Tissue Qiagen kit following the tissue protocol. Hematopoietic samples from the patient (whole blood, buffy coat, plasma, bone marrow) and PBMCs from family members and donors were also extracted with the DNeasy Blood and Tissue Qiagen kit following the blood protocol (buffy coat) or the cell protocol (rest of samples). DNA was quantified with a Thermo Qubit and quality checked with an Agilent TapeStation Genomic Tape.

### Duplex sequencing and variant calling

Libraries for duplex sequencing were prepared using TwinStrand kits (V1 chemistry) with 500ng input DNA. Libraries were captured with 3 different panel combinations depending on the sample type (**Supplemental Table S2**). All solid tissue samples were captured with the *TP53* panel, which includes the coding region and adjacent non-coding areas of the *TP53* gene (∼3,000 bp). The tumor tissues and a subset of normal tissues were captured with the *TP53* panel and a mutagenesis panel (TwinStrand’s Human Mutagenesis Assay), which includes probes covering 20 human genome regions spanning across 20 chromosomes designed to be an unbiased representation of sequence contexts and covering no loci with known selective advantages (48,000 bp, (Axelsson et al. 2024)). All blood samples were captured with the mutagenesis panel and a CHIP panel that includes the full coding sequence or hotspots of 29 genes recurrently mutated in adult AML and CHIP (TwinStrand’s AML-29 Assay, ∼59,000 bp). The CHIP panel includes *TP53* probes. Briefly, samples were digested with enzymatic fragmentation, end-repaired, A-tailed, ligated to TS adapters, amplified, and captured by hybridization with the corresponding probe panels. Two rounds of capture were performed to increase on-target reads. Then libraries were amplified, quantified, and pooled for sequencing on an Illumina NovaSeq 6000 using S4 flow cells. Raw reads were processed with the DNAnexus platform for duplex sequencing analysis with TwinStrand software version 3.20.1, which uses a variant caller based on VarDict (Lai et al. 2016). The pipeline produced duplex error-corrected VCF and depth by position files, which were used for downstream analyses.

### Mutation classification and pathogenicity

VCF file outputs were converted to MAF files for each sample with Vcf2Maf v1.6.22 using the Ensembl Variant Effect Predictor (VEP) tool version 112 to assign variant consequences. For each mutation, the variant allele frequency was determined by calculating the ratio of mutant duplex reads to the total duplex depth at the corresponding mutated position. Variants were discarded if they were in positions with depth lower than 1000 duplex reads, a no call fraction greater than 0.1, a variant allele frequency greater than 0.3 (corresponding to germline polymorphisms), or located in repeat regions defined using UCSC RepeatMasker and regions associated with known sequencing artifacts (bedfile provided at https://github.com/Colegrove/LiFraumeni_evolution/inputs/BEDs/LiFraumeni.mask.bed). To account for sequencing artifacts from potentially problematic regions, variants were discarded if they appeared in three or more samples and the DNAnexus-hosted pipeline annotated them as having 8 or more mismatches in the duplex read containing the mutation, suggesting a consistent pattern of mismatches. To account for potential SNP cross-contamination, variants at frequency greater than 0.3 in any sample were excluded from all other samples. Variants located within coding exons or extending up to two nucleotides into adjacent splice sites and indels that overlapped coding regions were considered coding variants. Mutation frequencies (MF) were calculated for the mutagenesis panel, CHIP coding mutations, *TP53* coding mutations, and *TP53* non-coding mutations as the number of unique mutations in each of those genomic regions divided by the total number of duplex nucleotides sequenced in those regions. We also calculated coding MF for each of the genes in the CHIP panel. Mutation burden (MB) was calculated for coding genes within the CHIP panel as the number of mutant duplex bases divided by the total sequencing depth across all coding positions in aggregate. All postprocessing analyses unless noted otherwise were performed with RStudio version 2025.09.1 using R version 4.5.1 and packages as indicated in **Supplemental Table S12**. Variants in *TP53* were assigned a pathogenic annotation and score based on AlphaMissense (Cheng et al. 2023) and mutation type. For missense variants, pathogenicity was annotated using AlphaMissense, which assigns a pathogenicity score between 0 and 1 such that values >0.564 are considered Likely Pathogenic, ≥0.34 but ≤0.564 are considered Ambiguous, and scores <0.34 are Likely Benign. Nonsense variants, indels, splice mutations, and multiple nucleotide substitution variants were considered Likely Pathogenic and assigned a score of 1. Synonymous mutations were assigned Likely Benign and a score of 0. Large clones were defined as mutations supported by more than one mutant duplex read, as each duplex read represents an independent DNA molecule.

### Testing for selection

To assess evidence for selection on somatic mutations, we calculated gene-specific *d*_N_/*d*_S_ ratios for LFS and non-LFS groups within CHIP panel genes. Synonymous and nonsynonymous SNVs were aggregated by gene and LFS status. Nonsynonymous mutations included missense, nonsense, and splice region mutations. The number of possible synonymous and nonsynonymous mutations for each gene was determined using release 115 of the Ensembl Variant Effect Predictor (VEP) web tool within the coding regions of the sequenced gene area. For mutations with multiple VEP annotated consequences, the most severe consequence as ranked by Ensembl was used. To enable *d*_N_/*d*_S_ calculations even in the absence of synonymous mutations, for each gene tested we added a pseudocount equal to the expected proportion of non-synonymous mutations to the numerator and the expected proportion of synonymous mutations to the denominator. For each gene and group, we resampled observed mutations with replacement 5,000 times to generate a bootstrapped *d*_N_/*d*_S_ distribution. For the *TP53 d*_N_/*d*_S_ analysis within LFS01’s tissues, samples were aggregated into broader anatomical categories (blood, solid tissue, sun-exposed skin, and cancer) to ensure sufficient sample sizes (**Supplemental Table S11**).

Phasing of *TP53* somatic mutations relative to the germline p.R181H allele was performed at the duplex consensus sequence-level using BAM files in Python version 3.12.3 with the pysam package v0.22.1. Somatic SNVs and indels located within 150 base pairs of the germline p.R181H site (hg38 chr17:7675070) were selected as candidate variants for phasing, corresponding approximately to the maximum duplex read length. If at least one duplex read spanned both the variant and the p.R181H site, we used the base identity of the p.R181H site to assign the variant to the LFS carrier or non-carrier *TP53* copy. Otherwise, the variant was designated as unphased. For variants containing more than one spanning duplex read, all duplex reads that could be phased were in agreement on the phase.

To contextualize patterns of mutation recurrence and see if mutations in the blood of all subjects and within the tissues of LFS01 overlapped with established recurrently mutated sites such as hotspots, we compared our observed *TP53* mutation distribution to reference data from the Catalogue of Somatic Mutations in Cancer (COSMIC) database v102 (GRCh38). COSMIC mutation data including missense, nonsense, and synonymous substitutions were downloaded as counts per amino acid mutation across all tumor types. We grouped entries by amino acid position and summed counts to generate total codon mutation frequencies. These frequencies were used to create reference lollipop plots aligned to the *TP53* SNVs across all LFS01 samples or across all subject blood samples.

### Regression analysis

First, to examine the relationships between coding mutation frequency (MF), mutagenesis MF, or mutation burden (MB) and age, we fit linear regression models. For each subject, we calculated coding and mutagenesis MF as the number of unique coding or mutagenesis mutations across all genes in the CHIP panel divided by the total number of coding or mutagenesis nucleotides sequenced. In addition, we calculated coding MB as the number of mutant coding bases divided by the total number of coding nucleotides sequenced. Linear models in the form Y∼ 1+X were fit separately for coding and mutagenesis MF or MB (Y) against age in years (X) using ordinary least squares regression implemented though the lm() function in R.

To further account for confounding variables, we performed multiple linear regression. The number of unique CHIP coding or mutagenesis mutations per 10 million duplex base pairs sequenced in the respective target region and the number of CHIP mutant coding bases per 10 million coding duplex base pairs sequenced were modeled separately with age (in decades), LFS status, and CTx status as covariates. We fit models in the form Y∼ 1 +age+LFS+CTx where LFS and CTx were treated as binary covariates using Gaussian generalized linear models with an identity link function as implemented in glm() in R. Analyses were also performed separately for each gene.

### Mutational spectrum and signature analysis

For a signature agnostic comparison between mutational spectra, we used AMSD v0.3.0 (Hart et al. 2026) performed using the “mean” option between p.R181H carrier and non-carrier groups and excluding samples with previous exposure to chemotherapy.

Mutational signatures were extracted to evaluate mutagenic processes in p.R181H carrier and non-carrier individuals. Only solid tissues sequenced with *TP53*+MUT panels were included in signature analysis and all hematopoietic samples. A separate signature extraction was performed for solid tissue samples and for hematopoietic samples. In both instances, single base substitutions were initially classified by their 96 trinucleotide contexts for pyrimidine-mutated base pairs using SigProfilerMatrixGenerator v.1.3.3. Mutational signatures were then extracted in each case using a Bayesian Hierarchical Dirichlet Process (mSigHDP, (Liu et al. 2023)). Tissue samples were pre-annotated as either ‘non-cancerous’ or ‘malignant,’ to facilitate hierarchical clustering of samples to enhance the accuracy of signature extraction (Liu et al. 2023). Extracted signatures from hematopoietic samples were decomposed using a recently published reference signature database for blood and chemotherapy-associated mutational signatures (Mitchell et al. 2025; Islam et al. 2022). Four mutational signatures were identified across hematopoietic samples: SBS1, SBS5, SBS7a, and SBSG. Extracted mutational signatures from solid tissue samples were decomposed using SigProfiler Extractor v.1.1.21 with COSMIC v3.4 reference database (Islam et al. 2022). Seven mutational signatures were identified from solid tissue samples: SBS1, SBS5, SBS7a/b/d, and SBS17a/b. Although LFS01 had received 5-fluorouracil chemotherapy, the signature for this agent (SBSG) was not incorporated into tissue analysis due to its similarity with SBS17a and SBS17b, which are known to be involved in esophageal adenocarcinoma (Abbas et al. 2023).

## Data access

All data associated with this study are provided in the paper or the Supplemental Materials. Sequencing data from this study will be made available through the NCBI BioProject database (https://www.ncbi.nlm.nih.gov/bioproject) prior to publication.

## Code availability

The code to reproduce all analyses can be found at https://github.com/Colegrove/LiFraumeni_evolution.

## Acknowledgements

We deeply thank the patient, the family, and the donors that provided samples, without whom this research would not have been possible. We thank the Willed Body Program at the University of Washington School of Medicine, the gastrointestinal oncology team at Fred Hutchinson Cancer Center who cared for this patient, the University of Washington autopsy service team who performed the pan tissue collection, and the University of Washington Medical Scientist Training Program classes of 2020 and 2021 (CONJ 511: Genomic Dissection) who provided excellent discussion about the case. We also thank Lauren Santos, program coordinator under Dr. Dubard-Gault at Fred Hutchinson Cancer Center, who helped consent individuals for this project and E. Jonlin for regulatory support.

## Author contributions

JJS, MSH, MEDG, and RAR conceived and designed the study; HLC, HM, RR, AF, and RAR designed the analysis; MEDG, DAM, EQK, JJS, MSH, and RAR contributed to sample acquisition; THS, ZKN, FYL, EKS, JEH, and CCV processed samples and performed sequencing and initial data analysis; HLC, HM, BFK, AF, and RAR analyzed the data; HLC, HM, RR, AF, and RAR interpreted the data; HLC, AF, and RAR wrote the article; JJS, MSH, and RR reviewed the article and provided feedback. All authors read and approved the final version of the manuscript.

