## Supplemental_Figures for "Ultra-deep duplex sequencing reveals unique features of somatic evolution in the normal tissues of a family with Li-Fraumeni syndrome"

Supplemental Figure S1: Duplex depth comparison across genomic regions and samples.

Supplemental Figure S2: CHIP coding regression of MF and age by gene.

Supplemental Figure S3: CHIP coding mutational burden association with age, chemotherapy, and LFS.

Supplemental Figure S4: Samples containing SBS7a are enriched for CC>TT dinucleotides.

Supplemental Figure S5: Comparison of pathogenicity of blood mutations in CHIP/AML genes in subjects with and without TP53 p.R181H germline mutation.

Supplemental Figure S6: Loss of heterozygosity (LOH) in Li-Fraumeni patient samples based on p.R181H allele frequency.

Supplemental Figure S7: Concordance of *TP53* mutations identified in blood samples and tissues from LFS01.

Supplemental Figure S8: Size and tissue distribution of *TP53* mutations observed in three or more samples.

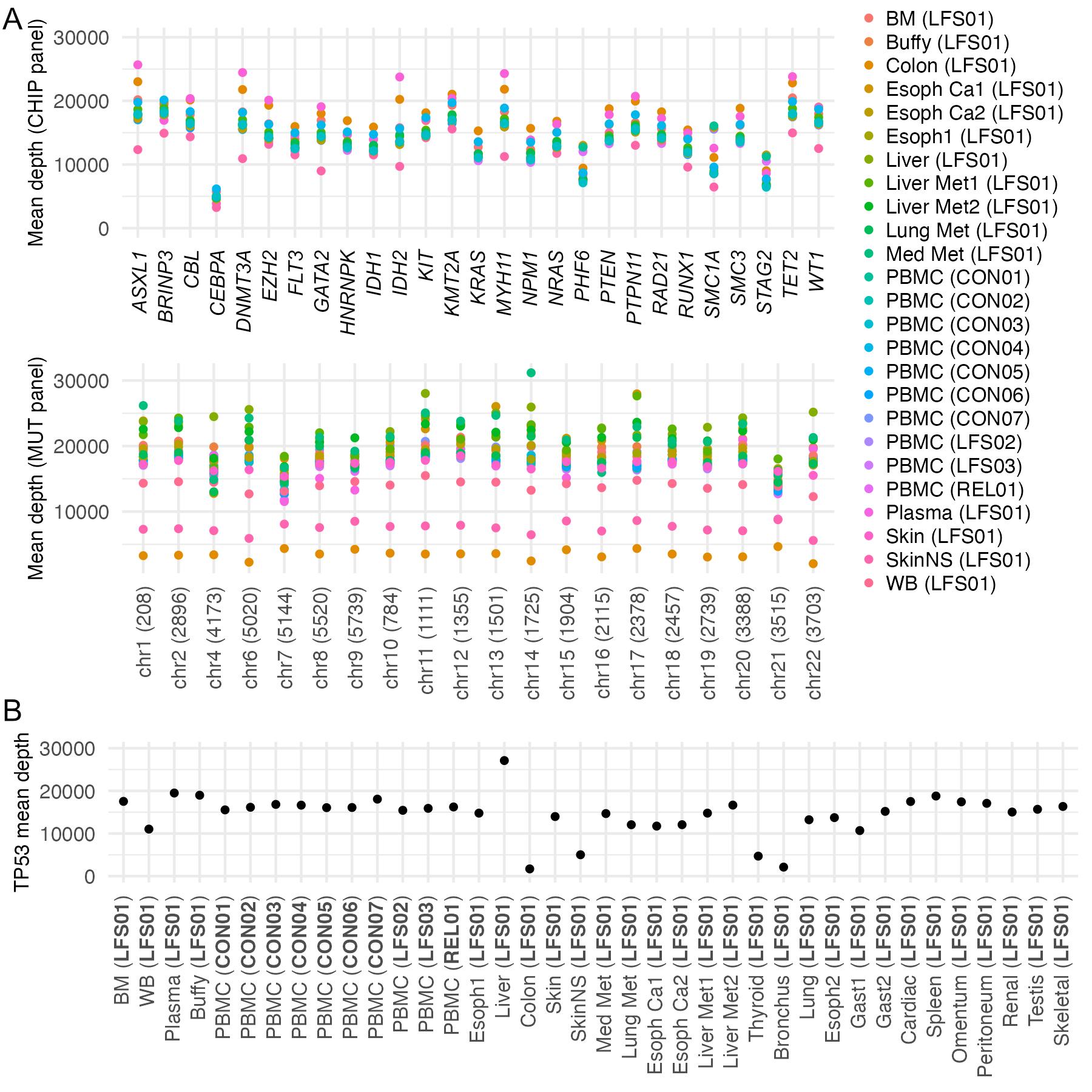

**Supplemental Fig. S1. Duplex depth comparison across genomic regions and samples.** (A) Mean duplex sequencing depth of CHIP panel genes and MUT panel regions colored by sample. (B) Mean duplex depth of TP53 by sample. BM: bone marrow; WB: whole blood; PBMC: peripheral blood mononuclear cells; Esoph: esophagus; Gast: gastric tissue; SkinNS: non-sun-exposed skin; Med: mediastinal; Met: metastasis; Ca: cancer.

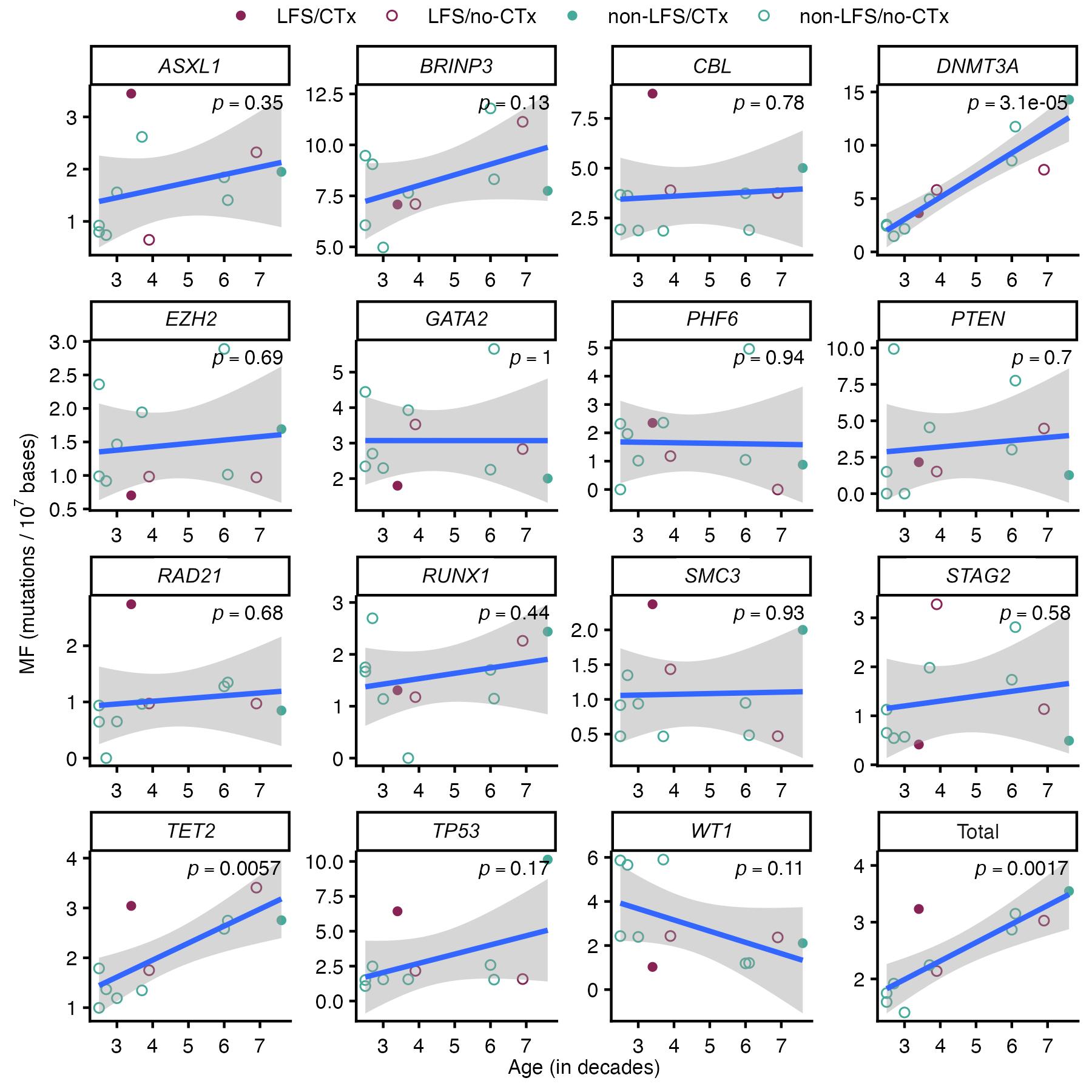

**Supplemental Fig. S2. CHIP coding regression of MF and age by gene.** Linear regressions of coding mutation frequencies (MF) in CHIP genes by age. Circles are colored by LFS (maroon) and non-LFS (teal) status and filled circles indicate chemotherapy (CTx) treatment. Age-associated baseline regression (blue line) is fit using all individuals and linear regression p-value is indicated. Grey areas indicate the confidence intervals of the regression line. Mutation frequency is measured as mutations per 10,000,000 bp sequenced.

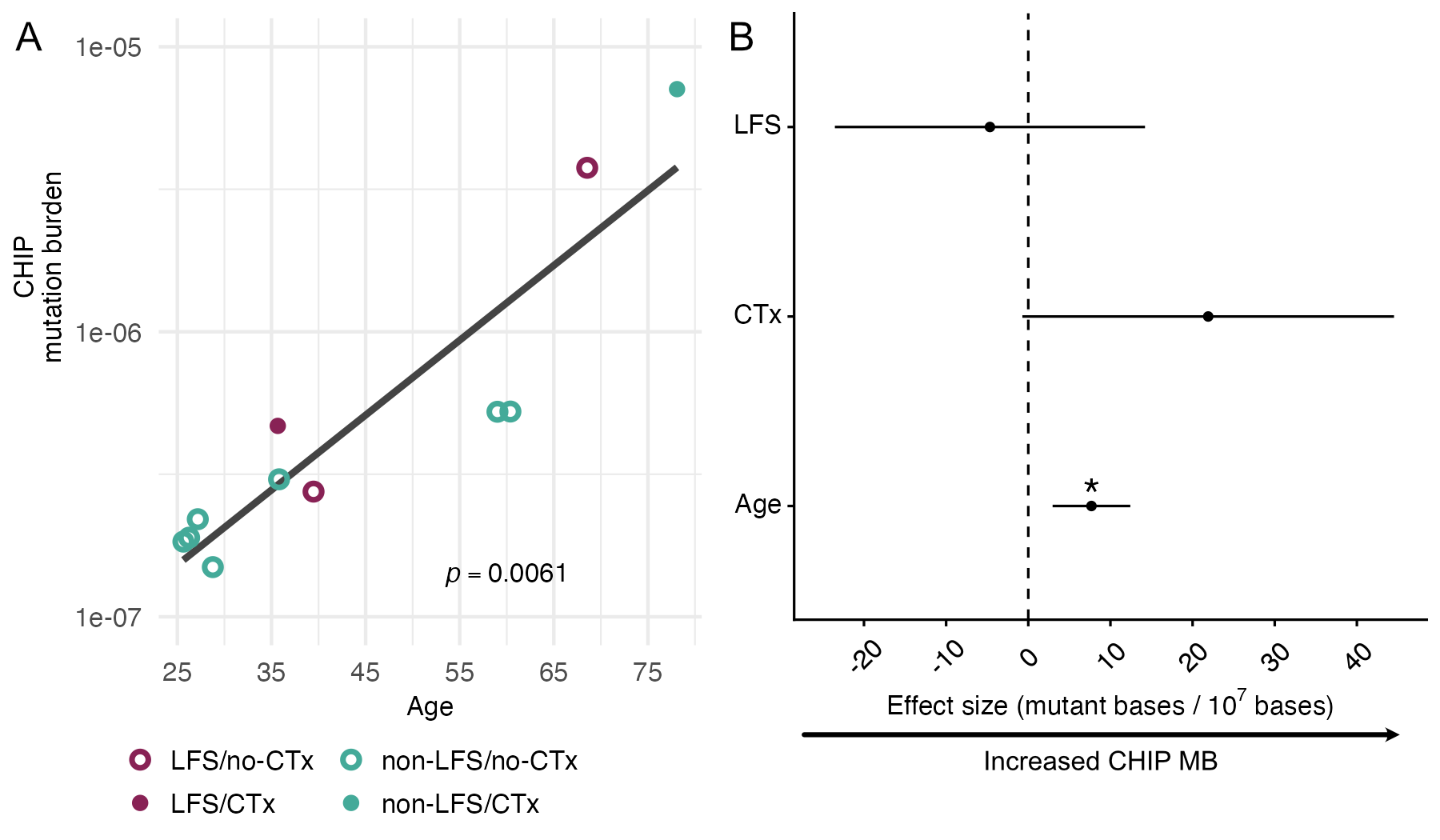

**Supplemental Fig. S3. CHIP coding mutational burden association with age, chemotherapy, and LFS.** (A) Linear regressions of coding CHIP mutation burden (MB) with age. Circles are colored by LFS (maroon) and non-LFS (teal) status and filled circles indicate chemotherapy (CTx) treatment. Age-associated baseline regression is fit using all individuals. (B) Multiple regression of coding CHIP MB using age (scaled by decades), LFS, and CTx status as covariates. MB was scaled per 10,000,000 bases. Asterisks indicate significance value (*p<0.05, **p<0.01) and error bars indicate 95% confidence intervals.

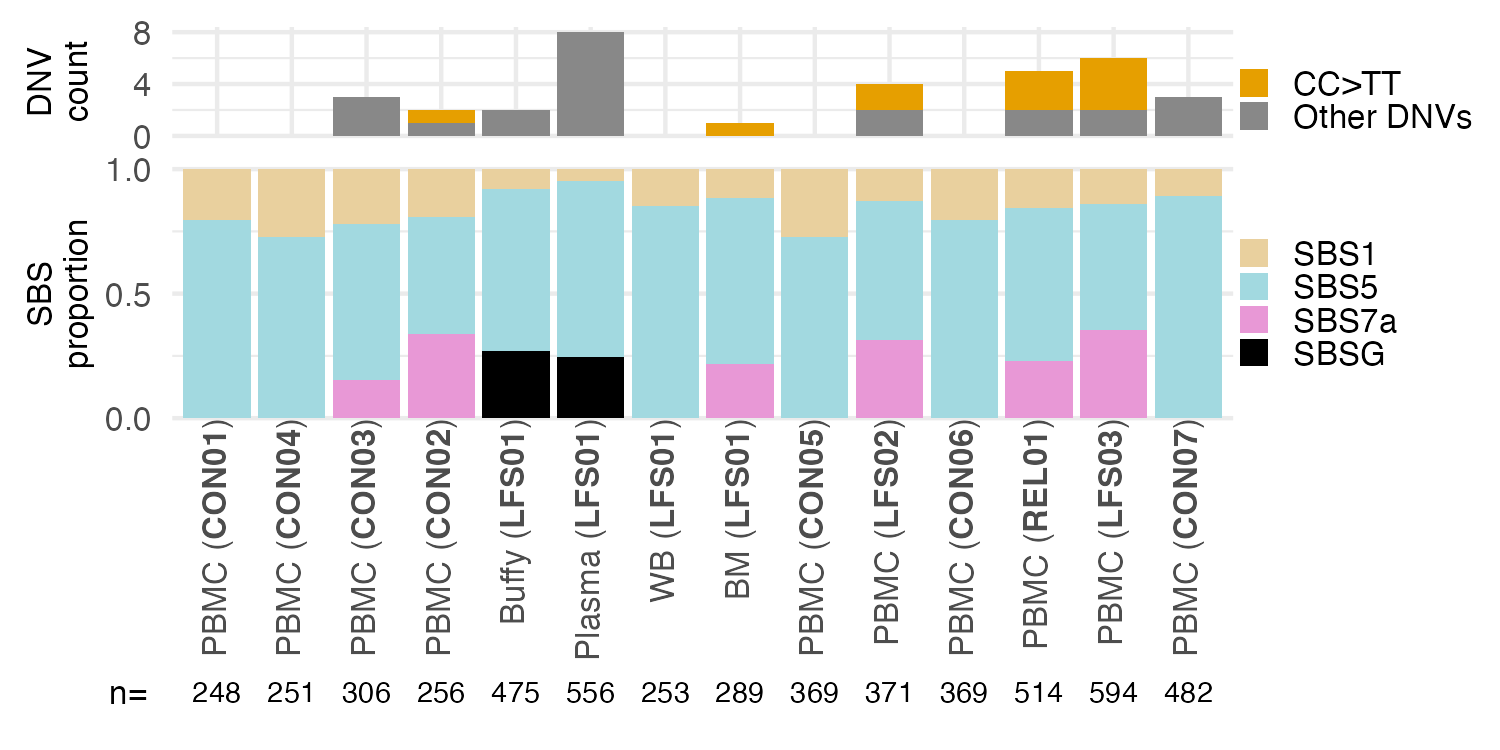

**Supplemental Fig. S4. Samples containing SBS7a are enriched for CC>TT dinucleotides.** Top: Bar plots showing dinucleotide variant (DNV) counts by subject and tissue with CC>TT (orange) and all other DNV classes (grey). Bottom: Proportion of mutations associated with SBS mutational signatures by subject and tissue. Total number of single nucleotide substitutions (n=) contributing to the signature decomposition for each sample are shown below the sample name.

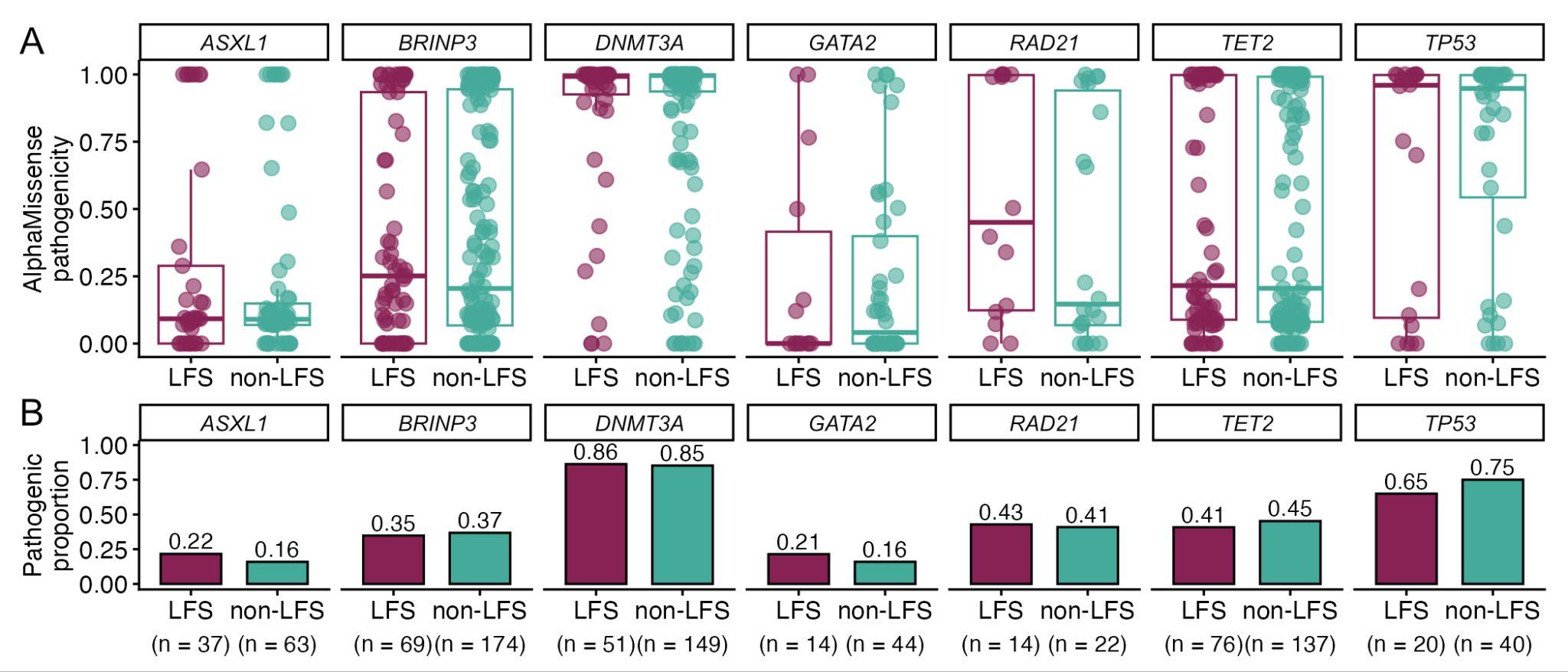

**Supplemental Fig. S5. Comparison of pathogenicity of blood mutations in CHIP/AML genes in subjects with and without TP53 p.R181H germline mutation.** (A) AlphaMissense pathogenicity scores of observed CHIP SNVs by germline p.R181H status (LFS: maroon; non-LFS: teal) within individual gene. Box plots indicate the median and interquartile range. (B) Percentage of pathogenic mutations from A. Pathogenicity is determined based on mutation type and AlphaMissense scores (see Methods).

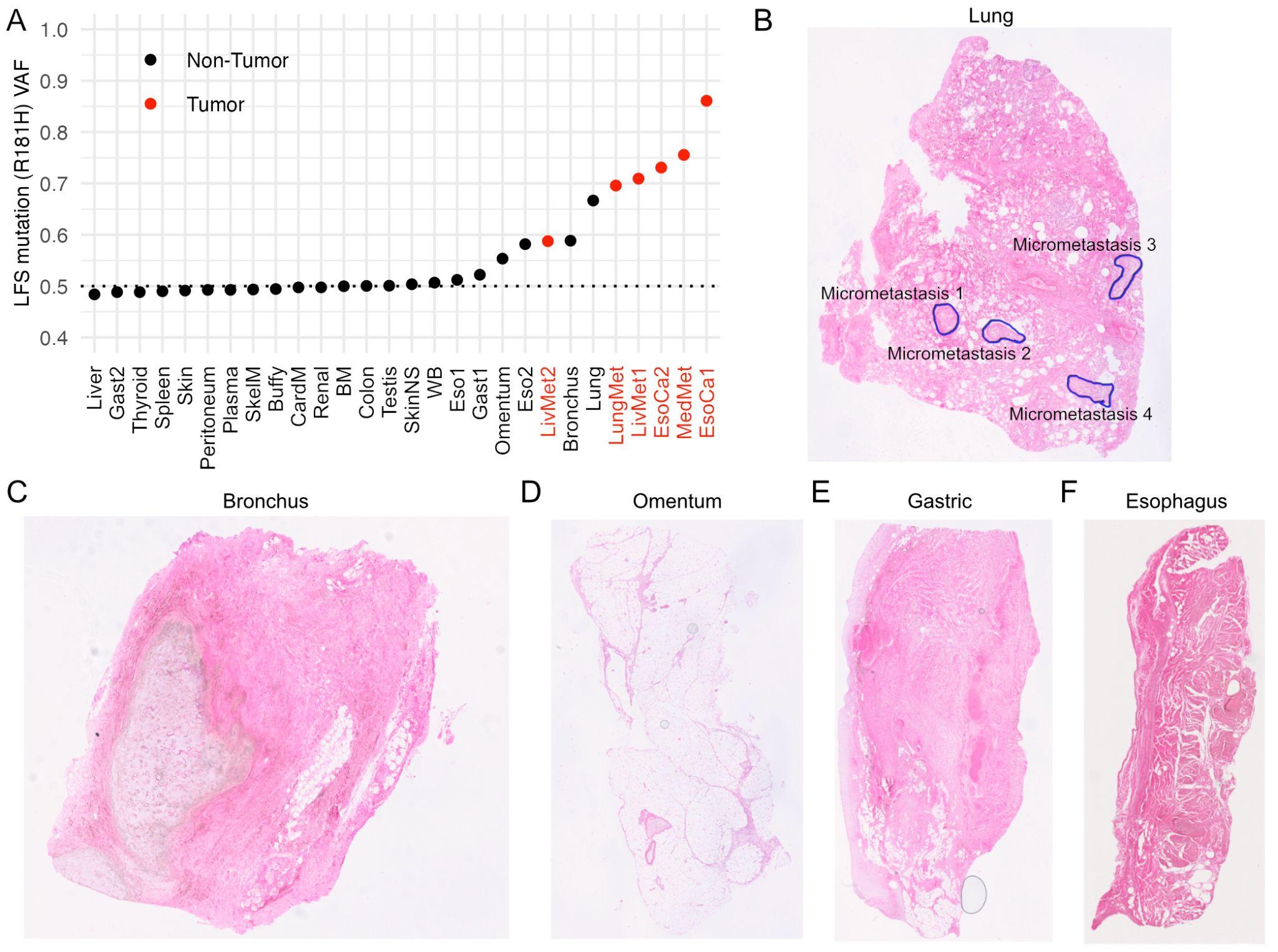

**Supplemental Fig. S6. Evaluation of loss of heterozygosity (LOH) and micrometastases in Li-Fraumeni patient samples.** (A) Variant allele frequency (VAF) of the p.R181H germline variant across LFS01 samples. Cancer samples indicated in red text. BM: bone marrow; WB: whole blood; PBMC: peripheral blood mononuclear cells; Esoph: esophagus; Gast: gastric tissue; SkinNS: non-sun-exposed skin; SkelM: skeletal muscle; Med: mediastinal; Met: metastasis; Ca: cancer. (B) Hematoxylin & eosin staining of lung biopsy in proximity to the one used for duplex-seq analysis. 3 micrometastases were identified. (C-F) Hematoxylin & easin staining of other normal tissues with p.R181H VAF>0.5. Micrometastases could not be identified in these sections, which were in proximity to the ones used for sequencing. Autopsy report, however, indicated fibrinous serositis and indurated omentum as well as fibrinous serositis and rare atypical cells in the gastric sample, suspicious of metastasis. (C) Brochus. (D) Omentum. (E) Gastric tissue. (F) Esophagus.

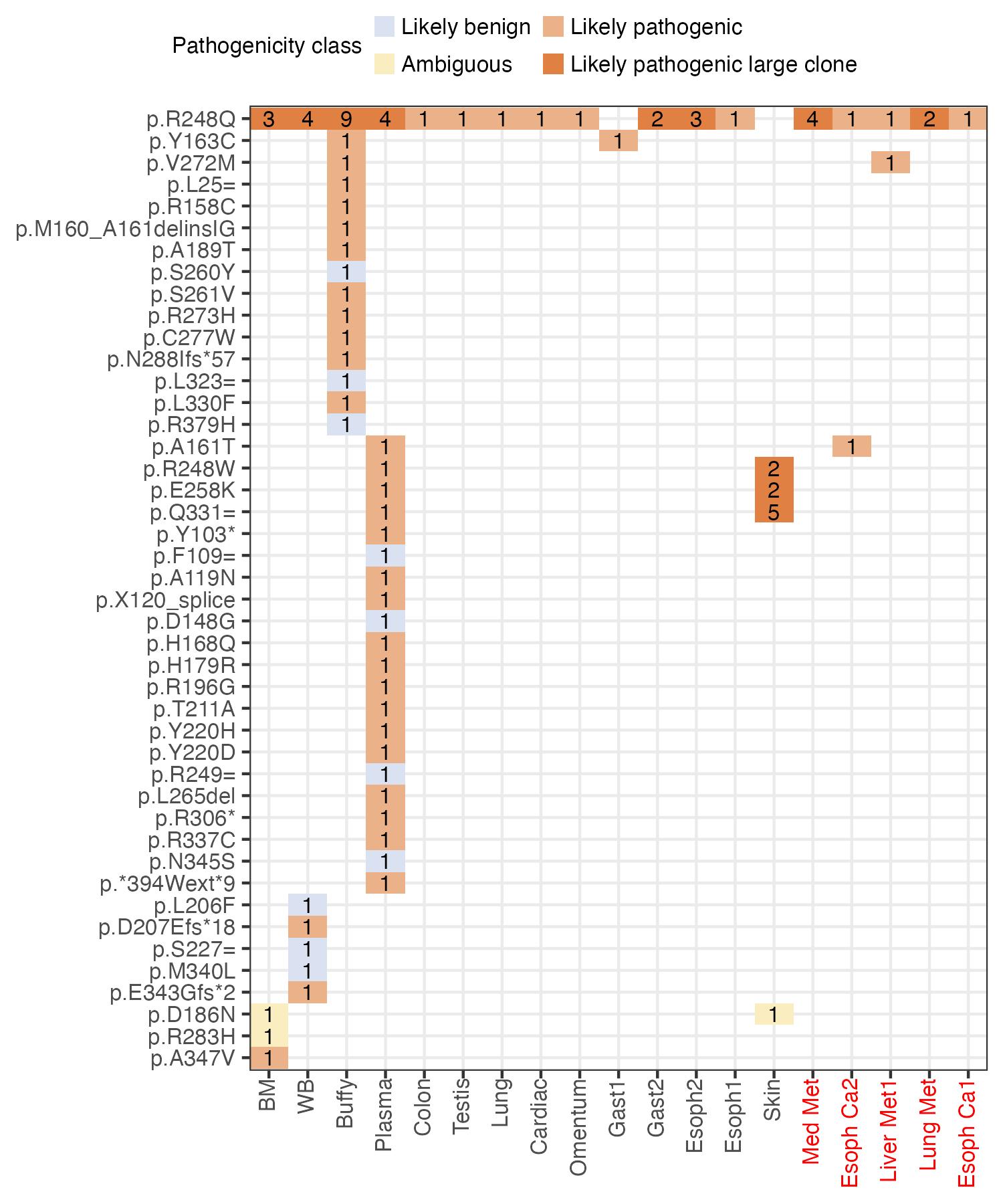
**Supplemental Fig. S7. Concordance of *TP53* mutations identified in blood samples and tissues from LFS01.** The plot includes all *TP53* mutations identified in bone marrow (BM), whole blood (WB), buffy coat (buffy), and plasma. Box color is determined by AlphaMissense pathogenicity score. Number inside of each box represents the number of duplex reads associated with that given mutation. Mutations with a mutant duplex read count greater than 1 are deemed a large clone and are colored a darker orange or blue depending on pathogenicity status. Cancer samples indicated in red text.

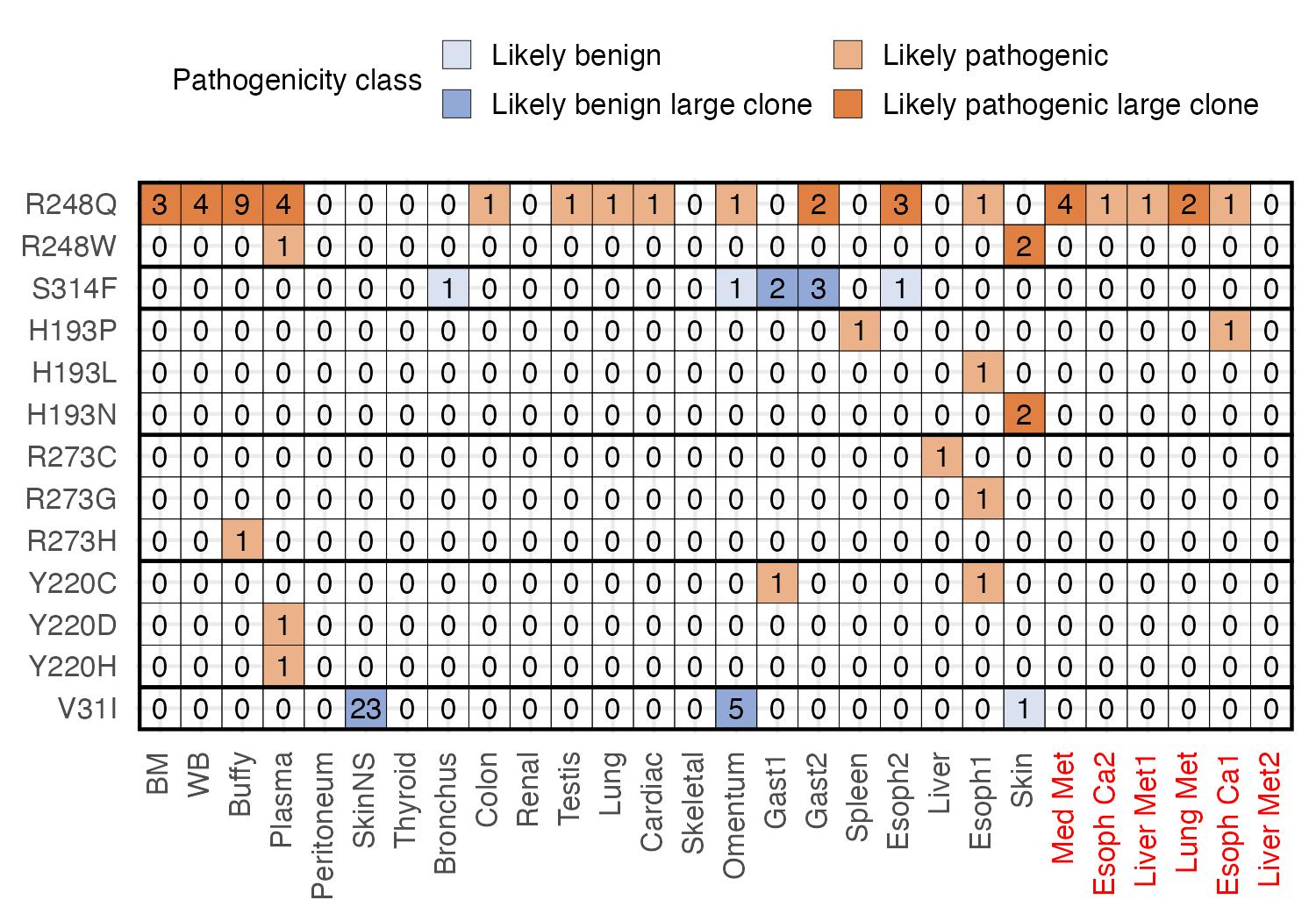

**Supplemental Fig. S8. Size and tissue distribution of *TP53* mutations in codons mutated in three or more samples.** Box color is determined by AlphaMissense pathogenicity score (likely pathogenic: orange; likely benign: blue). Number inside of each box represents the number of duplex reads associated with that given mutation. Mutations with a mutant read count greater than 1 are deemed a large clone and are colored a darker orange or blue depending on pathogenicity status. Cancer samples indicated in red text.
